# Measurement of hindered diffusion in complex geometries for high-speed single-molecule experiments

**DOI:** 10.1101/2020.07.04.187948

**Authors:** Tobias F. Bartsch, Camila M. Villasante, Ahmed Touré, Daniel M. Firester, Felicitas E. Hengel, Aaron Oswald, A. J. Hudspeth

## Abstract

In a high-speed single-molecule experiment, a protein is tethered between two substrates that are manipulated to exert force on the system. To avoid nonspecific interactions between the protein and nearby substrates, the protein is usually attached to the substrates through long, flexible linkers. This approach precludes measurements of mechanical properties with high spatial and temporal resolution, for rapidly exerted forces are dissipated into the linkers. Because mammalian hearing operates at frequencies reaching tens to hundreds of kilohertz, the mechanical processes that occur during transduction are of very short duration. Single-molecule experiments on the relevant proteins therefore cannot involve long tethers. We previously characterized the mechanical properties of protocadherin 15 (PCDH15), a protein essential for human hearing, by tethering an individual monomer through very short linkers between a probe bead held in an optical trap and a pedestal bead immobilized on a glass coverslip. Because the two confining surfaces were separated by only the length of the tethered protein, hydrodynamic coupling between those surfaces complicated the interpretation of the data. To facilitate our experiments, we characterize here the anisotropic and position-dependent diffusion coefficient of a probe in the presence of an effectively infinite wall, the coverslip, and of the immobile pedestal.

## Introduction

A protein under tension exhibits both entropic and enthalpic elasticity, a behavior that can be measured by observing the elongation of a single molecule while applying mechanical force. In such an experiment, the molecule is placed between two substrates, at least one of which is part of an elastic transducer through which forces can be delivered, for example an optically trapped, micrometer-sized bead. To avoid non-specific interactions between the protein and the substrates to which it is attached, the protein is usually secured through long, flexible DNA or PEG spacers. As a consequence, the fluctuations in the protein’s instantaneous position are filtered with a time constant of *γ*/*κ*, in which *γ* is the drag coefficient of the bead and *κ* is the total stiffness of the potential confining the bead, which comprises the spring constants of the optical trap, protein, and spacers. The position of the bead therefore reflects only the time-averaged end-to-end length of the protein. Moreover, information about the stiffness of the folded protein is concealed by the usually softer linker and often cannot be extracted from the measured force-extension relation. Resolving small structural changes and measuring the elasticity of folded proteins therefore remain challenging tasks that have recently been addressed through novel approaches. The fine structure of the energy landscape of DNA hairpins, for example, was measured with rigid DNA-origami spacers with a persistence length fiftyfold as great as the commonly used double-stranded DNA linkers^1^. Rigid spacers couple the motion of the protein’s ends tightly to the position of the bead, thereby increasing the bandwidth and precision of the experiment.

We developed a novel single-molecule assay that did not require long, flexible spacers. The protein was instead stretched directly between a diffusing probe—a 1 μm-diameter plastic bead to which force could be applied by optical tweezers—and an immobile glass pedestal—a 2 μm-diameter bead fixed to the coverslip. The protein’s ends were attached to the two beads through distinct, short, and relatively inelastic linkers.

This approach allowed us to characterize the equilibrium mechanics of protocadherin 15 (PCDH15), a protein whose properties implicate it as part of a molecular spring important for hearing^2^. Determining the entropic and enthalpic stiffness of the protein is crucial for our understanding of the molecular basis of mechanotransduction by the inner ear. Human ears can detect sounds at frequencies up to 20 kHz, and some bats and dolphins have a hearing range exceeding 200 kHz. The protein machinery that underlies hearing must therefore be capable of responding to very fast stimuli that likely produce mechanical responses far from thermal equilibrium. For two reasons, this high-frequency behavior has not been explored through single-molecule experiments. First, in the presence of flexible linker molecules, high-frequency force stimuli are largely filtered before they can elongate a protein of interest. Second, even in the absence of flexible linkers, the mechanical response of a protein is filtered owing to the drag on the bead and the stiffness of the optical potential that confines it. If these filtering effects are not too large compared to the time constant of the protein’s response, and if the drag on the bead is known, it is nevertheless possible to compensate for the filtering. In this study we characterize the anisotropic and position-dependent diffusion coefficient of a bead in our single-molecule assay in the presence of an effectively infinite wall, the coverslip, and of an immobile spherical obstacle, the pedestal. The results should facilitate analysis of high-speed single-molecule experiments relevant to auditory transduction.

## Results

### Correction of the position signal for light scattered by the pedestal

Determining the drag near a coverslip and pedestal requires high-precision measurement of the three-dimensional diffusion of a probe confined in a weak, position-sensing optical trap. The probe’s position can be estimated with sub-nanometer precision and microsecond temporal resolution by interfering the light scattered forward by the probe with the unscattered portion of the trapping beam on a quadrant photodiode^3^ (Fig. 1**A**). The diode’s difference signals are then linearly related to the probe’s position along the two axes perpendicular to the optical axis, and the signal summed over all four quadrants is proportional to the probe’s axial position.

**Figure 1.**
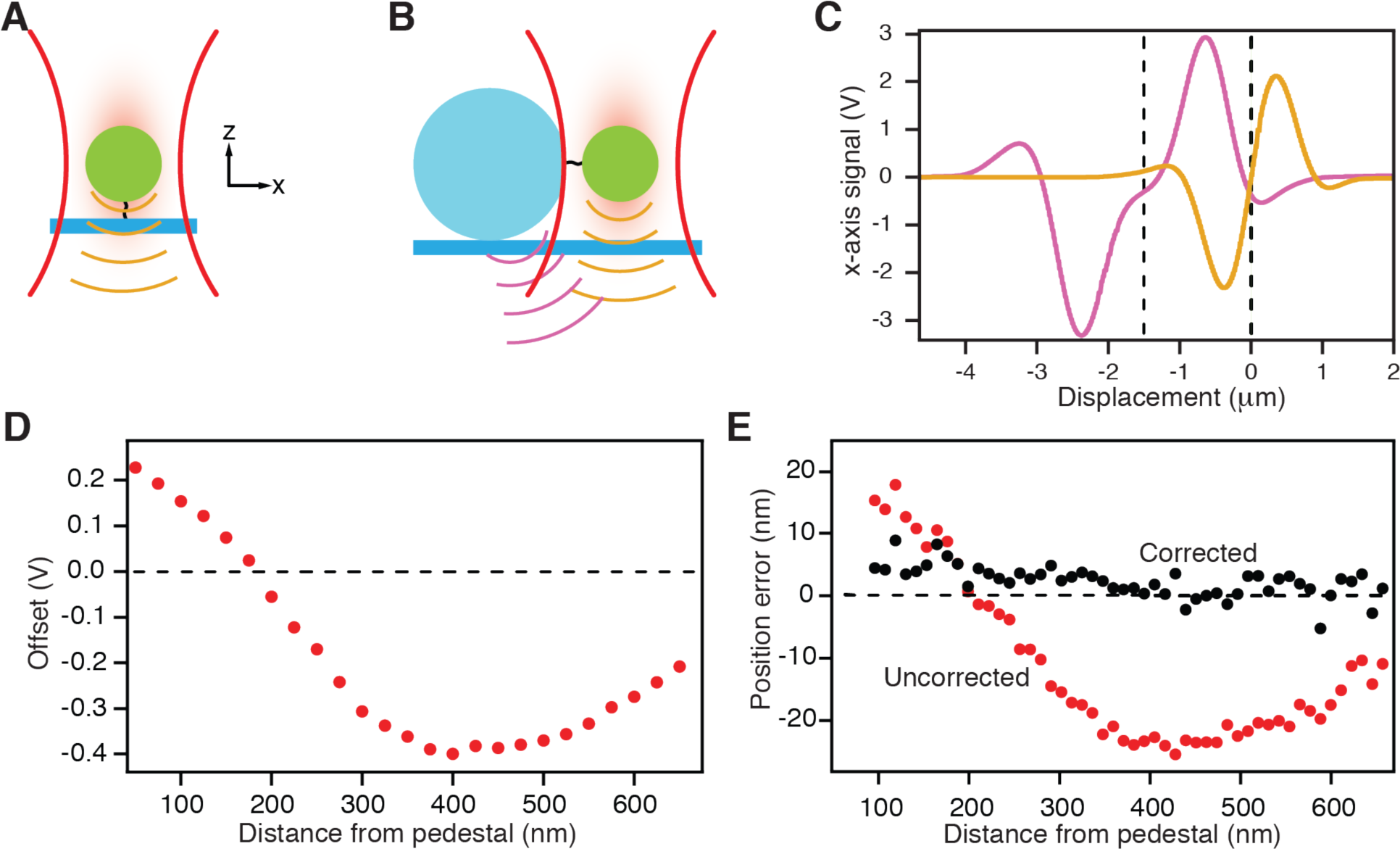
Apparatus and control experiments. **A**, When a probe (green) is held in an optical trap (red), its position can be measured along three axes by capturing transmitted and forward-scattered light (orange) on a quadrant photodiode. **B**, A stationary pedestal (blue), to which one end of a filamentous protein is attached, scatters a small fraction of the incident light (puce) and contaminates the desired signal for the probe. **C**, Control measurements show the signals as a function of offset position due to the probe alone (orange) and to the pedestal alone (puce). The two signals have been offset by 1.5 *μ*m (dashed lines) to simulate the configuration during an actual experiment. **D**, With the probe fixed in place, moving the pedestal nearby produces a spurious offset signal. The dashed line shows the reference signal for the probe far from the pedestal. **e**, The systematic error in position measurements owing to the pedestal is reduced by the compensation procedure to a few nanometers.

When the probe and pedestal are in close proximity—as is the case in single-molecule experiments without long linkers—the position-sensing beam is scattered not only by the probe, but also by the pedestal (Fig. 1**B**). Although this effect complicates estimation of the probe’s position, the diode’s total signal *S*_*total*_ can be approximated to first order as the sum of two independent signals^4,5^: the signal *S*_*pedestal*_ owing to the pedestal in the absence of the probe and the signal *S*_*probe*_ owing to the probe in the absence of the pedestal:

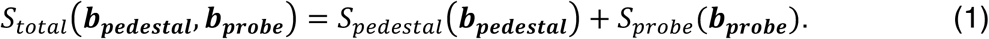

Here the vectors ***b*** represent the position coordinates of the probe and pedestal, which are the displacements of the respective objects from their positions when the photodiode’s output is zero. The offset *S*_*pedestal*_ is sensitive to the precise value of the distance ***b*_*pedestal*_** as well as to the shape of the pedestal itself, and must therefore be determined at the beginning of each experiment.

In a typical experiment, the pedestal is fixed at least 1.5 μm from the focal spot of the position-sensing beam, a distance determined by the radii of the pedestal and probe. The probe’s diffusion is confined by the beam’s trapping potential and is centered on the focal spot. The pedestal’s signal thus constitutes a constant offset added to the probe’s signal. If the magnitude of this offset is known, it can be subtracted from the total signal to yield the signal of the probe alone^2^.

To visualize the contributions of the two independent signals, we independently recorded the signals for displacements of the probe and the pedestal, then displayed them offset by 1.5 μm relative to one another (Fig. 1**C**). This procedure reflected the case in which the probe was at the center of the position-sensing optical trap, defined as *x* = 0, and just touched the pedestal. If the signal *S*_*probe*_ was held constant by fixing the probe’s displacement ***b*_*probe*_** from the focus of the position-sensing trap, then the offset could be determined by monitoring how the measured total signal *S*_*total*_ changed as the pedestal was brought progressively closer to the focal spot. Holding ***b*_*probe*_** constant by means of a second optical trap that strongly confined the probe at a displacement of 100 nm with respect to the focus of the position-sensing beam, we then recorded the total detector signal while the pedestal was so distant that its signal was negligible (*S*_*pedestal*_ ≈ 0). This signal served as a reference. As we moved the pedestal toward the focal spot of the position-sensing beam while keeping the probe confined at a constant position with the second trap, the deviation in *S*_*total*_ represented the signal *S*_*pedestal*_ owing to the pedestal (Fig. 1**D**).

In order to demonstrate that we could successfully correct for the influence of the pedestal, we next used the stimulus trap to hold the probe at the center of the position-sensing trap (*x* = 0). We recorded the photodiode’s total signal and recovered the position of the probe by subtracting the offset caused by the pedestal. The position signal after compensation was nearly zero (Fig. 1**E**). If the offset correction was not performed and the total signal on the detector was calibrated without subtraction of the pedestal’s influence, a significant systematic position error arose that depended sensitively on the distance between the pedestal and the center of the position-sensing optical trap. All the data presented in the remainder of this work were corrected by this means.

### Localization of the pedestal’s surface by thermal-noise imaging

Before assessing the hydrodynamic drag near a pedestal, it was necessary to localize the pedestal’s surface. We accomplished this by the super-resolution technique of thermal-noise imaging^6^. The spatial probability density of a probe diffusing in a weak optical trap was a three-dimensional Gaussian distribution with an ovoid iso-probability surface (Fig. 2**A**). When a pedestal intersected the optical trap, a portion of its volume became inaccessible to the probe’s diffusion: the forbidden volume in the probe’s spatial probability density then provided a negative image of the pedestal (Fig. 2**B**). We computed a line profile along the *x*-axis through the probe’s spatial probability density and converted the result by Boltzmann statistics to an energy landscape (Fig. 2**E**). We defined the wall of infinite energy as the impenetrable boundary of the pedestal and set *x* = 0 at this location.

**Figure 2.**
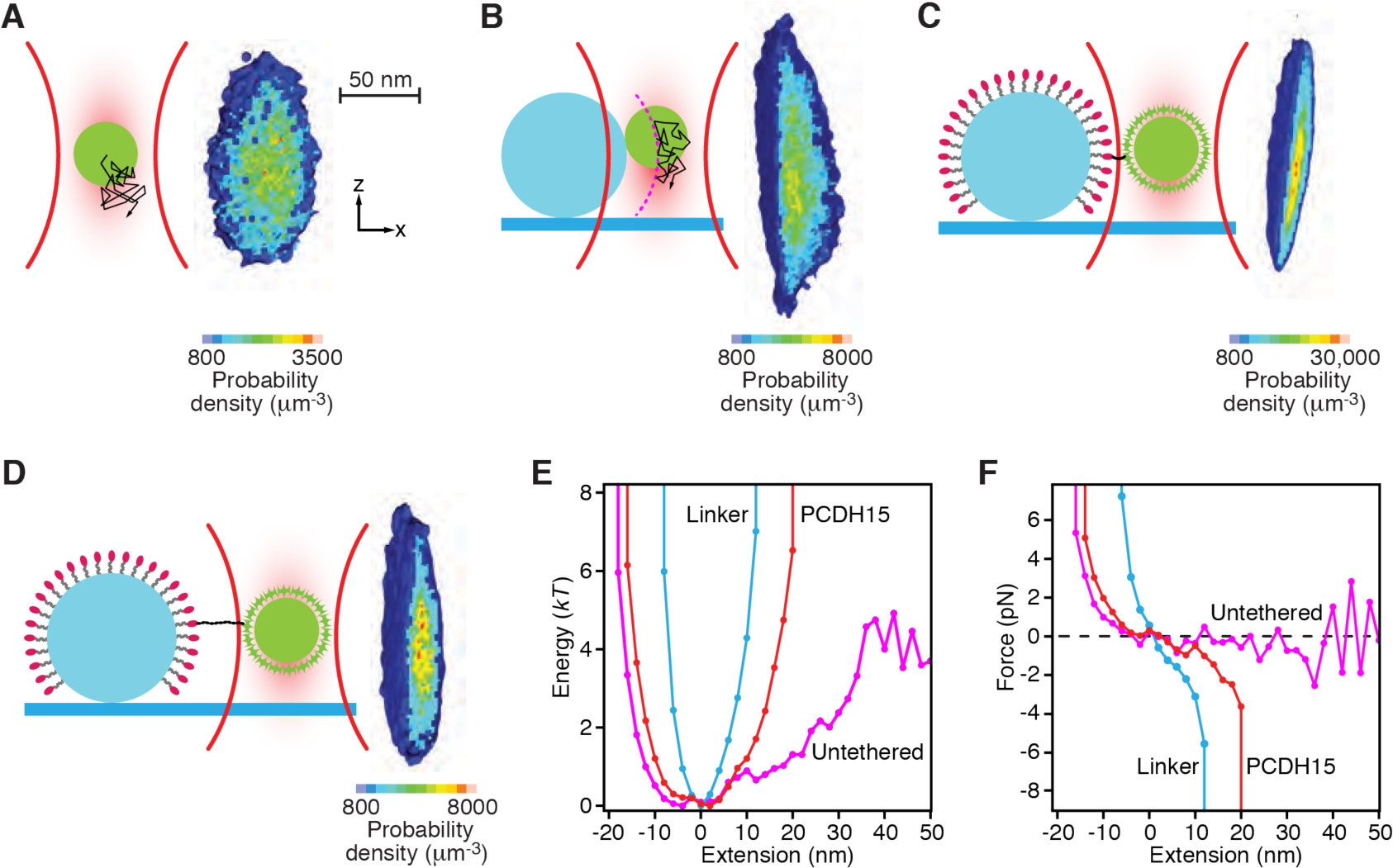
Determination of the pedestal’s position and trap’s strength. **A**, A schematic diagram (left) portrays the thermal diffusion of the probe in an optical trap. A section through the experimentally measured three-dimensional probability density (right) reveals the positions explored by the diffusing probe. The distribution is roughly symmetrical along the *x*- and *y*-axes, but elongated owing to weaker trapping along the z-axis. Note the discrepancy in scale: the density distribution is magnified about 25X in comparison to the 1 μm probe. The arrangements, definitions of axes, and spatial scales are identical in the following three panels. **B**, When the probe is brought into contact with the fixed pedestal, its diffusion is restricted. Flattening of the experimental probability density demarcates the surface of the pedestal. **C**, When the probe is affixed to the PEG-coated pedestal by a short linker, the linker further restricts diffusion of the probe. **D**, In an actual experiment, the probe is attached to the pedestal by a PCDH15 monomer. The protein’s extensibility allows the probe to explore a larger volume of space. **E**, The experimentally determined probability distributions reflect the energy of the system for the probe at various positions. **F**, The slopes of the displacement-energy relations in panel **E** represent the forces exerted on the probe by the optical trap and tethers.

Diffusion was further restricted when the probe was attached to the pedestal by a short peptide that represented the concatenation of the two linkers used in an experiment to attach a PCDH15 monomer to the probe and pedestal (Fig. 2**C**). In an actual experiment, the monomer was attached at each end by one of the linker peptides (Fig. 2**D**). In both instances, the energy functions became steeper as the probe was confined both by the optical trap and by the tether (Fig. 2**E**). The slopes of the three energy functions defined the position-dependent forces exerted on the probe (Fig. 2**F**).

### Determination of local diffusion constants

When a bead diffuses close to a boundary, its mean squared displacement becomes anisotropic and declines in comparison to that in bulk solution. Such hindered diffusion can be described by a position-dependent and anisotropic diffusion constant. Local diffusion constants have previously been measured by positioning an optically trapped bead at different distances from a boundary and computing the bead’s mean squared displacement^7^ or by inferring the drag from the power spectral density of the bead’s motion^8,9,10^. These methods average the drag’s value over the spatial extent of the bead’s diffusion in the relatively small volume of strong optical trapping.We instead confined a probe’s motion by a weak optical trap within a larger trapping volume of 160 nm x 140 nm x 253 nm in respectively the *x*-, *y*-, and *z*-directions. The beam profile was Gaussian along each axis, and this volume represented three standard deviations in each direction from the center of the beam. We then subdivided the trapping volume into voxels with edge lengths of 5 nm and computed the probe’s mean squared displacement independently within each voxel for a time lag of 150 μs (Fig. 3**A**)^11,12^. We then made use of the fact that, for each voxel, the slope relating the mean squared displacement along each axis to the time lag is twice the probe’s local diffusion constant along that axis. Although the resulting three-dimensional spatial map of diffusion constants was limited in spatial extent by the width of the trapping volume, larger volumes could be explored by displacing the optical trap in steps smaller than the width of the trapping volume and recording partially overlapping diffusion maps that were subsequently fused.

**Figure 3.**
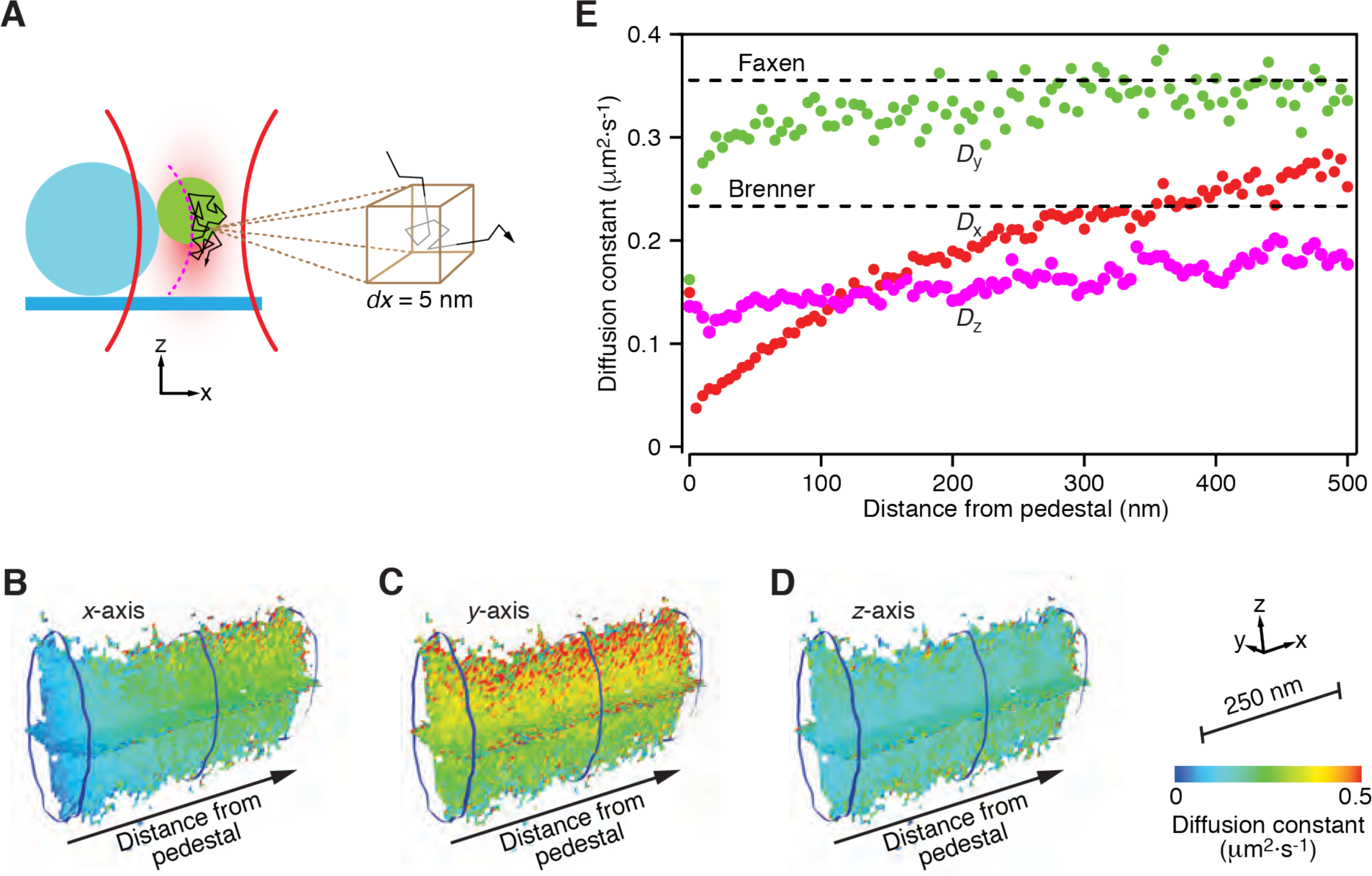
Measurement of local diffusion constants. **A**, A schematic diagram depicts measurement of the mean squared displacement for a probe centered in a 5 nm voxel. **B**, A heat map shows local diffusion constants along the *x*-axis as a function of the probe’s position along each of the three indicated axes. The definition of axes, spatial scale, and calibration are shown at the bottom right of the figure and pertain to the next two panels as well. **C**, Another map portrays the local diffusion constants along the *y*-axis. **D**, A similar representation displays the local diffusion constants along the *z*-axis. **E**, The local diffusion constants along the three axes display strikingly different behaviors. The values *D*_x_ for motion along the *x*-axis decrease sharply as the probe approaches the pedestal. Diffusion along the *y*-axis, parallel to the coverslip, yields values *D*_y_ relatively insensitive to position. The diffusion constants *D*_z_ for movement along the *z*-axis are reduced by proximity to the coverslip. These data are identical to those along the *x*-axes of the three preceding hear maps.

In our single-molecule assay of PCDH15 molecules, the protein was stretched along the *x*-axis. Because we were therefore mainly interested in how the associated diffusion constant *D*_x_ changed with extension from the pedestal, we moved the optical trap along that axis in 100 nm steps and determined the diffusion constant at each position (Fig. 3**B**). Although the focal spot of the optical trap remained fixed during each measurement, the trap was weak enough that the probe could diffuse along all three axes with respect to that point. We also computed the diffusion constants for motion along the *y*-axis, tangential to the pedestal but at a fixed height above the coverslip (*D*_y_, Fig. 3**C**) as well as those along the *z*-axis, tangential to the pedestal but perpendicular to the coverslip (*D*_z_, Fig. 3**D**). These results were determined for a probe maintained at a distance of 500 nm from the coverslip, so that the average *z*-position of the probe corresponded to the equator of the pedestal (Fig 3**E**).

Assuming that the coverslip acted as an infinite wall to which the probe’s diffusion coupled, we computed the diffusion constants expected in the absence of a pedestal for movements parallel and perpendicular to the coverslip as a function of the separation distance between the probe and coverslip. For positions far from the pedestal, we expected *D*_x_ and *D*_y_—the diffusion constants parallel to the coverslip—to approximate the value computed by Faxen’s law, whereas *D*_z_—the diffusion constant normal to the coverslip—was predicted to follow Brenner’s law^13^. As expected, the local diffusion constants progressively approached the analytical values with increasing distance between the probe and pedestal. The coupling of the probe’s diffusion to the pedestal extended beyond 500 nm along the x-axis, well in excess of the two tangential couplings, 35 nm for the *y*-axis and 445 nm for the *z*-axis. The ranges of the coupling were determined by the distances at which the experimental values approached within 15 % of the theoretical values. The diffusion constant along the *x*-axis declined by half upon extension from a separation distance of 50 nm to 200 nm, a range of particular interest for single-molecule experiments on the proteins that underly auditory sensation.

## Discussion

Precise characterization of the mechanical properties of a protein in a single-molecule experiment is dependent on accounting for factors that affect the measurements taken. In our experiments on the mechanical properties of PCDH15, a micrometer-sized probe serves as both a proxy for the position of the protein and the substrate through which forces are delivered to the protein. It is accordingly essential that measurements of the probe reflect the true mechanical response of the protein under study. The use of short, inelastic linkers reduces the filtering of the protein’s instantaneous position. Even for relatively short linkers, however, accurately measuring the effect of force on a protein requires compensation for the effects of hydrodynamic drag on the probe.

The use of short linkers introduces the additional complication that proximity of the pedestal and coverslip distorts the position signal of the probe. Here we presented a technique to measure and compensate for the influence of the pedestal. The offset measurements obtained by this means recovered the position of a probe at a known displacement from the center of the position-sensing trap with minimal error. In addition to aiding the accurate measurement of anisotropic and position-dependent drag, this technique demonstrates that the benefit of short linkers in single-molecule experiments needs not be limited by optical interaction with the substrates.

We then characterized the drag near the coverslip and near the pedestal. Using these results, we could compensate for the drag to which the probe is subjected. Our data show that the restricted diffusion of the probe when close to the pedestal is non-negligible in all directions. Of particular relevance to single-molecule experiments is the considerable restriction of diffusion along the *x*-axis, the direction of protein extension. The reduced diffusion constants associated with such motions significantly filter the mechanical response of a protein. Moreover, the diffusion constant in the direction of extension changes substantially over the range of distances relevant to single-molecule experiments involving auditory proteins.

Voxel size contributed to the resolution of our measurements, for smaller voxels permitted a more granular mapping of the local diffusion and drag. However, this benefit had to be balanced with the need to obtain sufficient data points from each voxel, the probability of which decreases as voxel size declines^12^. Another consideration for measurements of local diffusion constants was the choice of time lags at which to measure the mean squared displacement. This value plateaus beyond a characteristic autocorrelation time τ = *κ*/(*γk*_*B*_*T*) as a result of the probe’s confinement in the optical trap, resulting in a measured value smaller than that for a free particle. To capture the motion of the probe while it approximated free diffusion, the time lag accordingly had to be much smaller than this autocorrelation time.

A small time lag was also critical for another reason: the gradient force owing to the optical trap could result in drift. For a starting position far from the center of the trap, the gradient force causes the mean squared displacement to grow faster than free diffusion and therefore complicates measurements. The influences of the gradient force and of free diffusion can be compared by the relative mean-squared-displacement contribution^12^

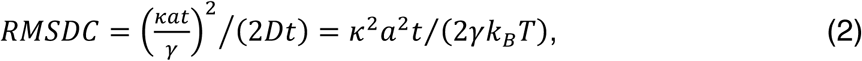

in which *κ* is the spring constant of the optical trap, which in our experiments was 4 μN·m^−1^ along the *x*- and *y*-axes and 1.2 μN·m^−1^ along the *z*-axis. *a* is the displacement from the trap’s center, *t* the time lag, *γ* the viscous drag coefficient of the probe given by Stokes’ law, and *D* the local diffusion constant. *k*_B_ and *T* are respectively the Boltzmann constant and thermodynamic temperature. For the ratio to remain small such that essentially free diffusion occurs, the time lag had to be set much smaller than the characteristic drift time *τ*_*D*_ = (2*γk*_*B*_*T*)/(*κ*^2^*a*^2^). Combining the two effects of optical trapping, the minimum of *τ* and *τ*_D_ determines the timescale at which the probe’s motion deviated from free diffusion. If *τ* < *τ*_D_, as was the case in our system for excursions of less than 150 nm from the trap’s center along any axis, then the influence of the gradient force on the mean squared displacement was negligible. If instead *τ* > *τ*_D_, then the measured value would have exceeded that of a freely diffusing particle for intermediate time lags.

## Methods

The photonic-force microscope used in these experiments was capable of measuring the position of a micrometer-sized probe bead with an integration time of 1 μs, sampled at 10^5^ s^−1^, with sub-nanometer precision. A weak optical trap was formed within the sample chamber by focusing a 1064 nm laser beam (Mephisto 500 mW, Coherent) with a high-numerical-aperture water-immersion objective lens (UPlanSApo 60xW, Olympus). In 81 experiments, the trap’s stiffness averaged 3.9 ± 1.0 μN·m^−1^ along the *x*-axis, 4.6 ± 1.1 μN·m^−1^ along the *y*-axis, and 1.1 ± 0.2 μN·m^−1^ along the *z*-axis (means ± standard deviations).

The three-dimensional position of the probe confined within the weak optical trap was obtained from the interference on a quadrant photodiode of light scattered forward from the probe with unscattered light. To hold the probe at a constant displacement from the center of weak optical trap, as was required for the correction owing to the pedestal, a stimulus trap was formed by an 852 nm laser (DL852-500, Crystalaser). The position of this relatively strong optical trap with respect to the weak trap was adjusted by means of a beam-steering lens in the beam path of the strong laser.

## Acknowledgments

TFB was supported by a Junior Fellow award from the Simons Foundation and CMV by MSTP grant T32GM007739 from the NIH. AT and DMF were supported by the F. M. Kirby Foundation. FEH received a Studienstiftung des Deutschen Volkes and AO a Medical Research Fellows Program grant from Howard Hughes Medical Institute, of which AJH is an Investigator.

## Authors’ Contributions

TFB, CMV, AT, FEH, and AO designed and conducted the experiments; TFB, CMV, AT, FEH, AO, and DMF analyzed the data; TFB, CMV, AT, FEH, AO, DMF, and AJH wrote the paper.

## References

1. Pfitzner, E. et al. Rigid DNA Beams for High-Resolution Single-Molecule Mechanics. Angew. Chem. Int. Ed. 52, 7766–7771 (2013).

2. Bartsch, T. F. et al. Elasticity of individual protocadherin 15 molecules implicates tip links as the gating springs for hearing. Proc. Natl. Acad. Sci. U. S. A. 116, 11048–11056 (2019).

3. Pralle, A., Prummer, M., Florin, E. L., Stelzer, E. H. & Hörber, J. K. Three-dimensional high-resolution particle tracking for optical tweezers by forward scattered light. Microsc. Res. Tech. 44, 378–386 (1999).

4. Bartsch, T. F., Kochanczyk, M. D., Lissek, E. N., Lange, J. R. & Florin, E.-L. Nanoscopic imaging of thick heterogeneous soft-matter structures in aqueous solution. Nat. Commun. 7, 1–9 (2016).

5. Seitz, P. C., Stelzer, E. H. K. & Rohrbach, A. Interferometric tracking of optically trapped probes behind structured surfaces: a phase correction method. Appl. Opt. 45, 7309 (2006).

6. Bartsch, T. F. et al. Detecting sequential bond formation using three-dimensional thermal fluctuation analysis. Chemphyschem Eur. J. Chem. Phys. Phys. Chem. 10, 1541–1547 (2009).

7. Pralle, A., Florin, E.-L., Stelzer, E. H. K. & Hörber, J. K. H. Local viscosity probed by photonic force microscopy. Appl. Phys. Mater. Sci. Process. 66, S71–S73 (1998).

8. Jannasch, A., Mahamdeh, M. & Schäffer, E. Inertial Effects of a Small Brownian Particle Cause a Colored Power Spectral Density of Thermal Noise. Phys. Rev. Lett. 107, 228301 (2011).

9. Franosch, T. et al. Resonances arising from hydrodynamic memory in Brownian motion. Nature 478, 85–88 (2011).

10. Schäffer, E., Nørrelykke, S. F. & Howard, J. Surface Forces and Drag Coefficients of Microspheres near a Plane Surface Measured with Optical Tweezers. Langmuir 23, 3654–3665 (2007).

11. Hsu, Y.-H. & Pralle, A. Note: Three-dimensional linearization of optical trap position detection for precise high speed diffusion measurements. Rev. Sci. Instrum. 85, (2014).

12. Tischer, C., Pralle, A. & Florin, E.-L. Determination and Correction of Position Detection Nonlinearity in Single Particle Tracking and Three-Dimensional Scanning Probe Microscopy. Microsc. Microanal. 10, 425–434 (2004).

13. Happel, J. & Brenner, H. Low Reynolds number hydrodynamics. vol. 1 (Springer Netherlands, 1981).

